# NRP2 as an emerging angiogenic player; promoting endothelial cell adhesion and migration by regulating recycling of α5 integrin

**DOI:** 10.1101/744763

**Authors:** Abdullah AA Alghamdi, Christopher J Benwell, Samuel J Atkinson, Jordi Lambert, Stephen D Robinson

## Abstract

Angiogenesis relies on the ability of endothelial cells (ECs) to migrate over the extracellular matrix via integrin receptors to respond to an angiogenic stimulus. Of the two neuropilin (NRP) orthologs to be identified, both have been reported to be expressed on normal blood and lymphatic ECs, and to play roles in the formation of blood and lymphatic vascular networks during angiogenesis. Whilst the role of NRP1 and its interactions with integrins during angiogenesis has been widely studied, the role of NRP2 in ECs is poorly understood. Here we demonstrate that NRP2 promotes Rac-1 mediated EC adhesion and migration over fibronectin (FN) matrices in a mechanistically distinct fashion to NRP1, showing no dependence on β3 integrin (ITGB3) expression, or VEGF stimulation. Furthermore, we highlight evidence of a regulatory crosstalk between NRP2 and α5 integrin (ITGA5) in ECs, with NRP2 depletion eliciting an upregulation of ITGA5 expression and disruptions in ITGA5 cellular organisation. Finally, we propose a mechanism whereby NRP2 promotes ITGA5 recycling in ECs; NRP2 depleted ECs were found to exhibit reduced levels of total ITGA5 subunit recycling compared to wild-type (WT) ECs. Our findings expose NRP2 as a novel angiogenic player by promoting ITGA5-mediated EC adhesion and migration on FN.

## Introduction

Neuropilins (NRPs) are single non-tyrosine kinase receptors belonging to a family of type I transmembrane glycoproteins (MW ∼130–140 kDa) [1]. To date, two NRP orthologs have been identified in vertebrates, NRP1 and NRP2, both of which share a very similar domain structure and an overall 44% amino acid homology [2, 3]. Their expression and function have been identified in many cell types, including nerve cells, endothelial cells (ECs), epithelial cells, immune cells, osteoblasts and tumour cells [3, 4]. In ECs, it is believed that NRPs play an essential role in sprouting angiogenesis and lymphogenesis through the selective binding to members of the vascular endothelial growth factor (VEGF) family. Following this complex formation, NRPs function as co-receptors with VEGFRs to enhance the VEGF-induced activation of many intracellular pathways [5]. As such, the functions of NRP have been implicated in influencing cell adhesion, migration and permeability during angiogenesis, under both physiological and pathological conditions [6-8]. Studies have, however, provided evidence to suggest that NRPs can mediate ligand signalling independently of VEGFRs, in addition to regulating VEGFRs independently of VEGF binding. Based on early transgenic mouse studies in this field, it was originally speculated that NRP1 is mainly expressed on arteries, arterioles and capillaries, whereas NRP2 is expressed on veins, venules and lymphatic vessels [9, 10]. However, subsequent studies have revealed that both NRPs are expressed in normal blood and lymphatic endothelial cells, and both play essential roles in forming blood and lymphatic vasculature networks [11-13]. Despite this, investigations into elucidating the roles of endothelial NRPs during angiogenesis have, for the most part, focused on NRP1 [3]. With regard to NRP2, studies have instead focused on annotating a role in cancer cells, where its upregulation is consistent with cancer progression in a number of cell types (e.g. neuroblastomas [14], non-small cell lung carcinoma [NSCLC] [15], human prostate carcinoma, melanoma [4], lung cancer [15-17], myeloid leukaemia [18], breast cancer [19] and pancreatic cancer [20]). Interestingly, Favier *et al*. [21] recapitulated this upregulation of NRP2 observed in cancer cells by overexpressing NRP2 in human microvascular ECs (hMVECs), and found that cell survival in these ECs was significantly increased following stimulation with either VEGF-A- or VEGF-C. Furthermore, NRP2 knockdown significantly inhibited both VEGF-A- and VEGFC-induced migration, suggesting that NRP2 as a potential pharmacological target. Crucially, this work highlighted the importance of understanding the cross-talk between NRP2 and other receptors (mainly VEGF, integrins and plexins) in ECs, which will aid in the design of novel drugs to better control the mechanisms underlying angiogenesis and lymphangiogenesis in autoimmune diseases and tumour development [21]. We have previously described a link between NRP1 and the β3-integrin (ITGB3) subunit during VEGF-induced angiogenesis. We reported that complete loss of the Itgb3 gene enhanced EC permeability through the upregulation of VEGF-VEGFR2-ERK1/2 signalling [22], and that NRP1 and ERK1/2 expression was elevated in ITGB3-NULL ECs. Subsequent targeting of NRP1 in ITGB3-NULL mice revealed a significant inhibition of VEGF-induced angiogenesis compared to WT mice, indicating that the elevation of angiogenesis in the absence of the Itgb3 gene is dependent on NRP1 expression [8, 23]. In this study we aimed to investigate whether NRP2 shares a similar interaction with ITGB3 or other integrins in ECs.

## Results

### NRP2 function is not regulated by ITGB3 during VEGFR2-mediated signalling or migration over FN matrices

We previously showed the involvement of NRP1 during VEGF-stimulated angiogenesis to be dependent upon ITGB3 in ECs [8]. Due to the structural homology between NRP1 and NRP2 [3], we decided to first consider whether, like NRP1, NRP2’s function shares a dependency on ITGB3 during VEGF-mediated angiogenic responses in ECs. To investigate this, we isolated mouse lung microvascular endothelial cells (mLMECs) from both wild-type (WT) and ITGB3-heterozygous (β3HET) mice and immortalised them with polyoma-middle-T antigen (PyMT) by retroviral transduction. DNA was extracted from multiple immortalised lines, and analysed by PCR to confirm their genetic status as either WT or β3HET cells (**Suppl. Fig1A**). We subsequently confirmed the EC identity of each immortalised line by examining the expression of EC markers including VE-Cadherin [24, 25], in addition to quantifying the expression of ITGB3 in β3HET ECs. Each clone was confirmed to express canonical EC markers, and all β3HET ECs were confirmed to express approximately 50% ITGB3 compared to their WT counterparts (**Suppl. Fig1B-D**). We reported previously that in β3HET ECs, NRP1 expression is upregulated, and that NRP1 appeared only to play a role in post development angiogenesis when ITGB3 expression is reduced [8]. Western blot quantification of NRP2 expression between WT and β3HET protein lysates showed a similar elevation in NRP2 expression in β3HET ECs (**Fig1A**), suggesting the existence of a regulatory nexus between NRP2 and ITGB3, which we felt warranted further exploration.

**Figure 1.**
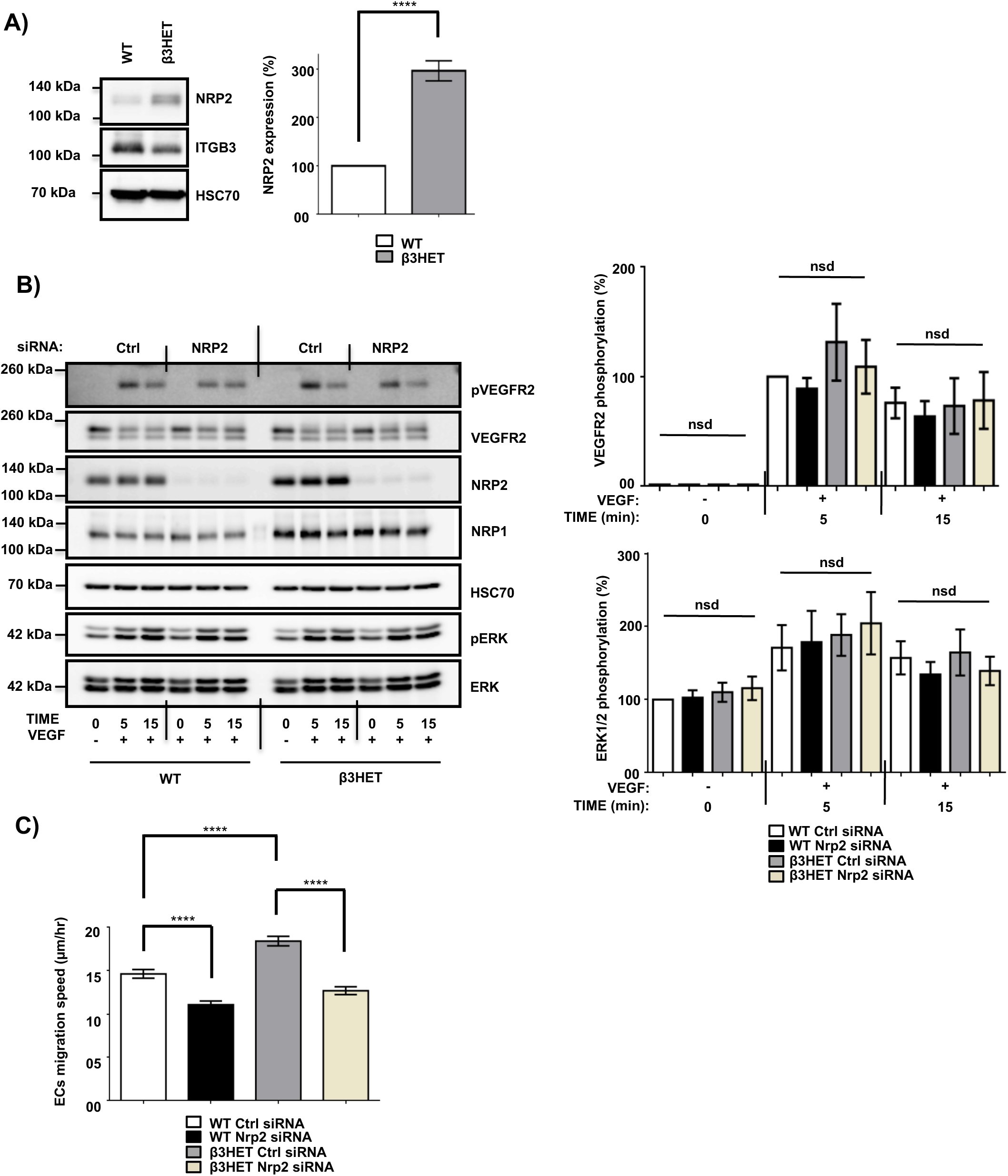
NRP-2 is not regulated by ITGB3 during VEGFR-2-mediated signalling or during migration over FN. (**A**): WT and β3HET ECs were plated onto FN and incubated for 48 hours at 37°C and 5% CO_2_. EC extracts were immunoblotted using antibodies to NRP-2, ITGB3 and HSC70. N=3 independent experiments, ****P < 0.0001. (**B** and **C**): siRNA-transfected ECs were plated onto FN and incubated for 48 hours at 37°C and 5% CO_2_. ECs were subsequently starved in serum-free media and stimulated with VEGF (30 ng/ml) for the indicated timepoints. EC extracts were immunoblotted using antibodies to phospho-VEGFR-2, VEGFR-2, NRP-2, NRP-1, HSC70, phospho-ERK and ERK. N=6 independent experiments, P > 0.05 (nsd). Left panel shows representative Western blot images, right panel shows densitometric analysis of band intensities normalised against HSC70 and obtained using ImageJ^TM^. Asterisks indicate statistical significance, nsd indicates no significant difference from unpaired two-tailed t-tests. (**C**): siRNA-transfected ECs were seeded onto FN and incubated for 3 hours at 37°C and 5% CO_2_. Fixed images were taken every 10 minutes for 15 hours at 37°C and 5% CO_2_ using an inverted Zeiss Axiovert microscope with one-phase contrast. Random migration speed was quantified using the ImageJ^TM^ plugin mTrackJ in µm/hour. N= 4 independent experiments, n≥30 ECs per experimental condition, ****P < 0.0001.

As NRP2 has been shown to regulate VEGF-induced signalling in both human lymphatic [26], and lymphatic microvascular ECs, we examined whether NRP2 regulates proangiogenic signalling responses to VEGF and if the effects are dependent on ITGB3. Using NRP2 specific siRNA (**Suppl. Fig1E**) versus a control siRNA transfected into our WT and β3HET ECs, we measured differences in VEGFR2 and ERK phosphorylation over a time course of 15 minutes. We observed only marginally attenuated VEGFR2 and ERK phosphorylation by Western blot analysis in response to NRP2 silencing and, importantly, saw no differences between WT and β3HET ECs (**Fig1B**), suggesting that ITGB3 does not regulate NRP2 dependent VEGF induced signalling in our cells.

To explore the possibility of a regulatory axis between NRP2 and ITGB3 further, we examined cellular migration. Angiogenesis relies on the ability of ECs to respond to angiogenic stimuli by migrating over an extracellular matrix (ECM). We have shown previously that NRP1 plays a role in promoting EC migration over fibronectin (FN) matrices, but only when ITGB3 levels are reduced [8]. Independent studies have also described NRP2 silencing to inhibit migration of human microvascular ECs [21] and human lymphatic ECs [26]. We therefore chose to examine the effects of NRP2 depletion on EC migration over FN in our mLMECs, and to determine whether any effect is dependent upon ITGB3 expression. To achieve this, control and NRP2 siRNA transfected WT and β3HET ECs were plated on FN and random migration speed was measured by time-lapse microscopy over 15 hours. As previously reported, we observed β3HET ECs migrate faster over a FN matrix than WT ECs [8]. However, unlike NRP1, whilst depletion of NRP2 significantly reduced EC migration speed, it did so independently of ITGB3 expression (**Fig1C**). We therefore conclude that ITGB3 does not regulate NRP2 function during these angiogenic processes.

### Depletion of NRP2 in ECs disrupts adhesion to FN matrices

Whilst the upregulation of NRP2 expression we observe in β3HET ECs suggests a regulatory crosstalk between NRP2 and ITGB3 (**Fig1A**), NRP2 regulated signalling and migration show no dependence on ITGB3. As NRP2 depletion significantly impaired the ability of ECs to migrate over FN (**Fig1C**), but not to undergo proliferation (**Suppl. Fig2A**) we next chose to directly examine the effect of NRP2 depletion on cell adhesion to FN. To do this we compared the relative number of cells adhered to 96-well plates pre-coated with FN for either 15 or 30 minutes. At both timepoints significantly fewer cells adhered to FN following NRP2 siRNA treatment compared to control siRNA treated ECs (**Fig2A**). Migration is dependent on the ability of cells to adhere to a matrix via the formation of FA complexes that continuously cycle through phases of assembly and disassembly. FAs comprise a module of recruited intracellular proteins, such as paxillin and integrins, that link to the actin cytoskeleton to mediate mechanical changes in the cell [27-30]. Whilst NRP2 depletion had no effect on total or phosphorylated levels of paxillin in our ECs (**Fig2B**), we did observe it to negatively impact the rate of focal adhesion (FA) turnover itself. FA assembly and disassembly were monitored in ECs treated with either control or NRP2 siRNAs, transfected with paxillin-GFP, by time-lapse microscopy for 30 minutes over a FN matrix. Silencing of NRP2 in mLMECs significantly reduced the rate of both FA assembly and disassembly compared to control siRNA treated cells (**Fig2C**). We also measured FA number and size distribution in ECs by immunolabelling for endogenous paxillin in cells that had adhered to FN for 90 minutes, a time that allows for mature FAs to form [31-33]. Despite no significant difference in average cell area (**Suppl. Fig2B**), FAs were significantly fewer in number and smaller in average size in NRP2 siRNA treated cells compared to control siRNA treated ECs (**Fig2D**), suggesting NRP2 depletion inhibits FA maturation. Finally, we considered whether NRP2 promotes EC adhesion and migration on FN by regulating Rac1 activation. Rac1 is a small Rho GTPase that mediates cell motility following integrin engagement, regulating leading edge cytoskeletal dynamics and promoting the formation of FAs [34-37]. Using a recombinant PBD-domain protein (PAK-1) fused to glutathione-magnetic beads, we captured the relative abundance of active GTPase-GTP in control and NRP2 siRNA treated ECs stimulated for 180 minutes on FN. Compared to our control siRNA treated ECs, NRP2 depleted ECs exhibited a significantly reduced level of active Rac1 (**Fig2E**), suggesting that NRP2 promotes EC adhesion and migration on FN by regulating Rac1 activation.

**Figure 2.**
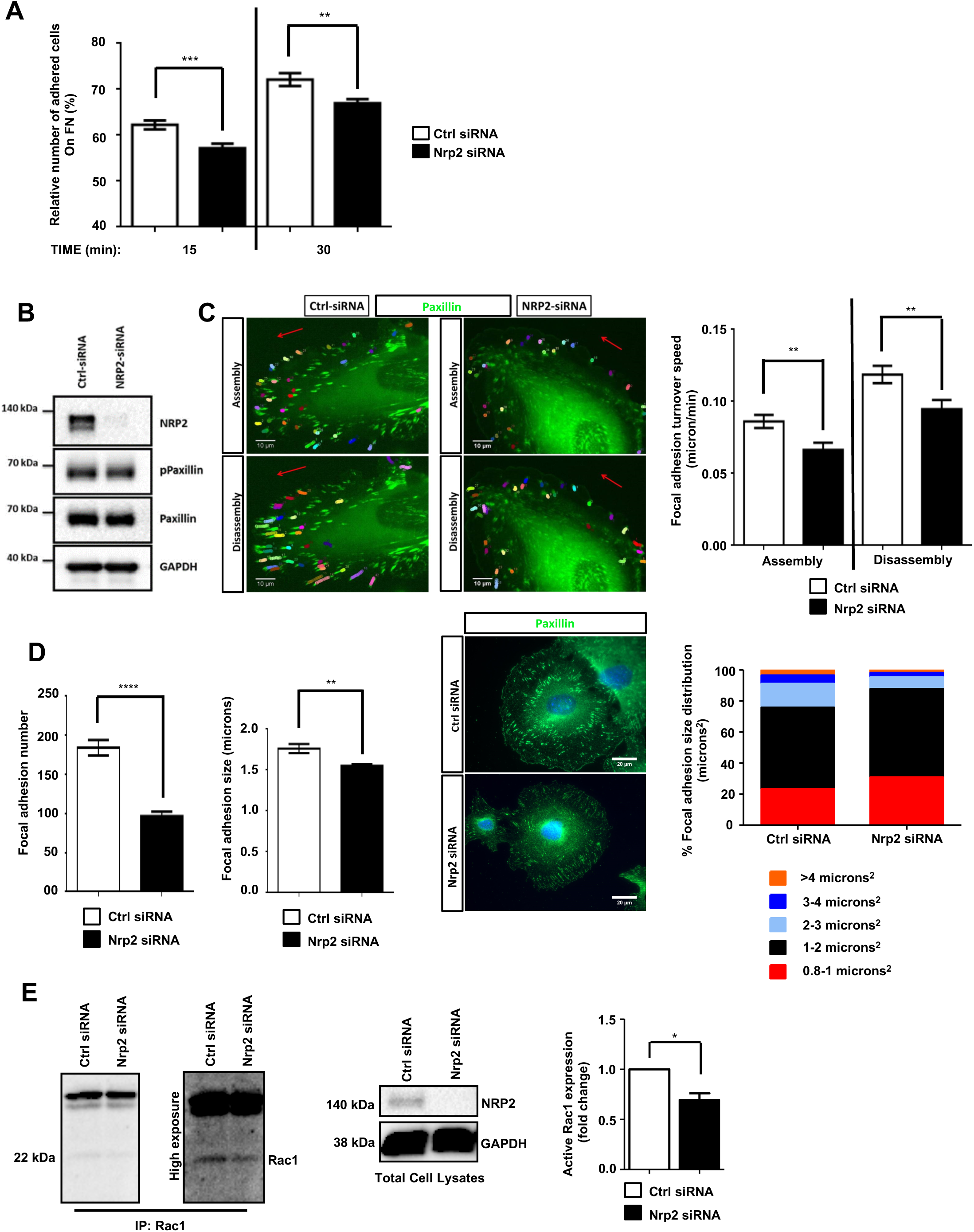
Depletion of NRP2 in ECs disrupts migration and adhesion to FN matrices by affecting Rac1 activation. (**A**): siRNA-transfected ECs were seeded onto FN coated 96-well plates pre-blocked 5% bovine serum albumin (BSA), and incubated for either 15 or 30 minutes at 37°C and 5% CO_2_. ECs were subsequently fixed in 4% PFA and stained with methylene blue. Absorbance was read at 630 nm. Data was normalised to the relative number of the total cells seeded in a 3-hour incubation control plate. N=3 independent experiments, ** P<0.01, *** P<0.001. Asterisks indicate statistical significance from unpaired two-tailed t-tests. (**B**) siRNA-transfected ECs were plated onto FN and incubated for 48 hours at 37°C and 5% CO_2_. EC lysates were immunoblotted using antibodies to NRP2, phospho-paxillin, paxillin and GAPDH. (**C**) siRNA-transfected ECs were double transfected with a GFP paxillin construct, seeded onto FN and incubated for 48 hours at 37°C and 5% CO_2_. ECs were fixed in a Ludin chamber and live imaged at 37°C and 5% CO_2_ using an inverted Axiovert microscope in which an individual cell was captured every one minute for 30 minutes (left panels). Focal adhesion (FA) assembly and disassembly speeds were analysed using the ImageJ^TM^ plugin mTrackJ in µm/min. In each respective bar, n≥100 FAs, ** P < 0.01 (right panel). (**D**) siRNA-transfected ECs were seeded onto FN and incubated for 90 minutes at 37°C and 5% CO_2_ before being fixed and stained for paxillin. Images were taken using a Zeiss AxioImager M2 microscope at 63x magnification. FA number and size was quantified using ImageJ^TM^ software as previously described by Lambert *et al*. [68]. A FA size lower detection limit was set at 0.8 microns. The centre panel shows representative images for fixed ECs transfected either with control or NRP-2 siRNA. Image quantification for FA number and size is shown in the left panel. Quantification performed on mean data from n≥25 ECs over N=3 independent experiments. % FA size distribution analysis is shown in the right panel on mean data from n≥25 ECs over N=3 independent experiments. Asterisks indicate statistical significance from an unpaired two-tailed t-test. (**E**) siRNA-transfected ECs were seeded onto FN and incubated for 180 minutes at 37°C and 5% CO_2_. EC extracts were immunoprecipitated by incubation with 10 µg Rac1 assay reagent (PAK-1 PBD magnetic beads) for 45 minutes at 4°C with gentle agitation. Immunoprecipitated complexes were subjected to Western blot analysis using antibodies against Rac1. Left panel: low and high exposure images showing active Rac1 levels in control and NRP2 siRNA transfected lysate. Middle panel: NRP2 depletion was confirmed by Western blot analysis using antibodies against NRP2 and GAPDH. Right panel: densitometric analysis of band intensities normalised against GAPDH and obtained using ImageJ^TM^. Asterisks indicate statistical significance from an unpaired two-tailed t-test.

### NRP2 regulates ITGA5 expression in ECs

Cells adhere to the ECM via heterodimeric integrin receptors, which are recruited to FA complexes during cell migration [38, 39] and which activate Rho family GTPases such as Rac1 [35, 36]. α5β1 integrin is the principle FN binding integrin in ECs [7, 29], and has been previously described as being upregulated during developmental angiogenesis to promote EC migration and survival [40, 41].Studies have also shown NRP1, through its cytoplasmic SEA motif, to specifically promote α5β1 integrin-mediated EC adhesion to FN matrices in a VEGF independent fashion [7, 42]. As NRP2 depletion significantly impaired mLMEC migration and adhesion on FN, and given the structural homology shared between NRP1 and NRP2, we considered whether a similar association exists between NRP2 and α5β1 integrin. First, we examined whether NRP2 knockdown regulated the expression of either integrin subunit. Western blot analysis revealed that siRNA-mediated silencing of NRP2 resulted in a significant upregulation of ITGA5 subunit expression in four different EC lines (**Fig3A**), whilst β1-integrin (ITGB1) expression remained unchanged (data not shown). Whilst endothelial ITGA5 specifically pairs with ITGB1 [39], ITGB1 can form heterodimers with α subunits 1 to 9 (with the exception of α7) [43, 44], suggesting α5β1-integrin behaviour can be studied by examining ITGA5 discretely (e.g. ITGB1 expression profiles are sum measures of its interactions with multiple α subunits present in the cell). Therefore, we chose to subsequently focus our attentions on ITGA5 and its interplay with NRP2.

**Figure 3.**
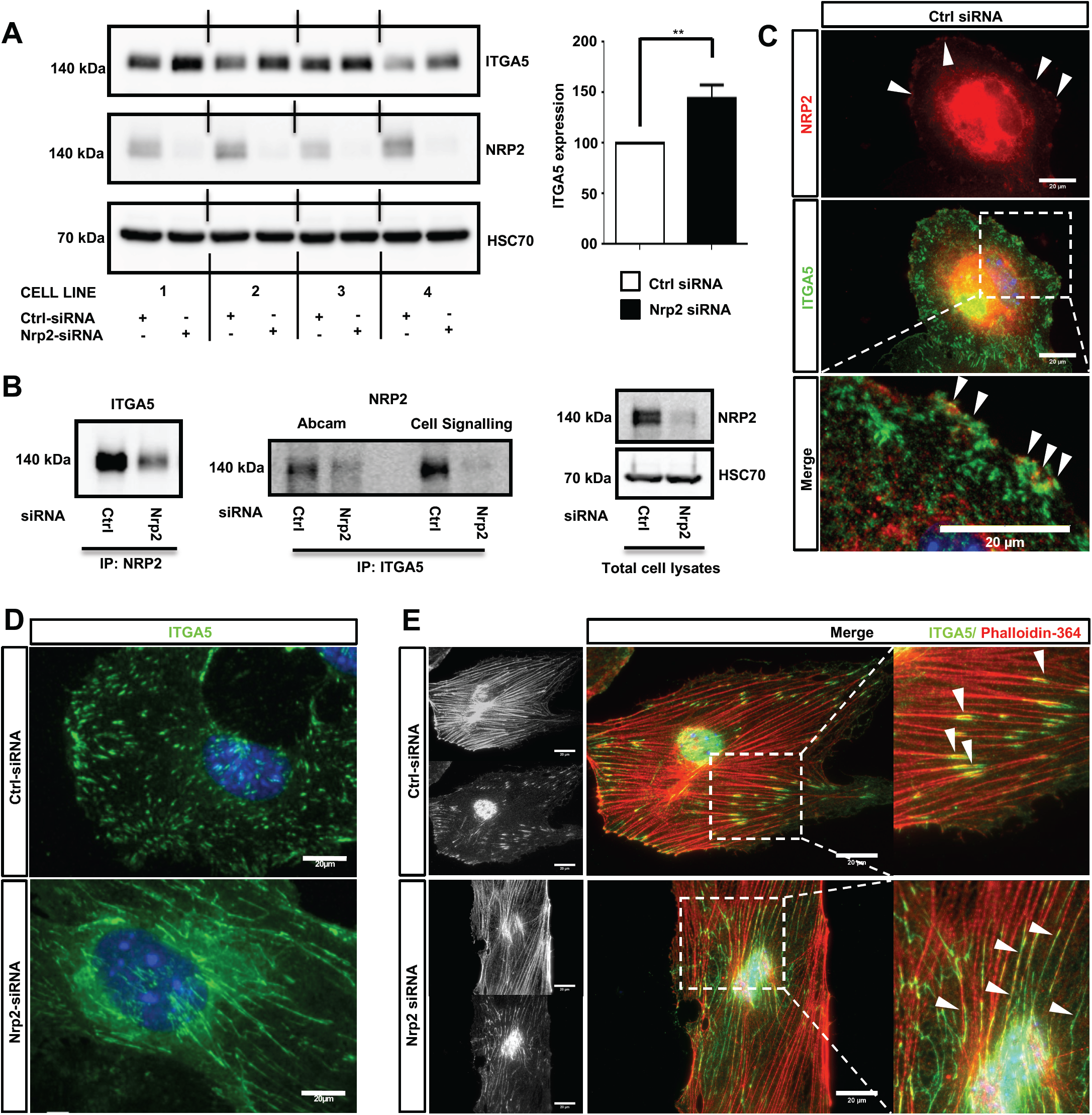
NRP-2 regulates ITGA5 expression in ECs. (**A**): siRNA-transfected ECs from four different clones were seeded onto FN and incubated for 48 hours. EC extracts were immunoblotted using antibodies to ITGA5, NRP-2 and HSC70. Left panel shows Western blot image, right panel shows densitometric analysis of ITGA5 band intensities normalised against HSC70 and obtained using ImageJ^TM^. ** P<0.01. (**B**): siRNA-transfected ECs were seeded onto FN and incubated for 48 hours. Left panel: EC extracts were immunoprecipitated by incubation with protein-G Dynabeads® coupled to a NRP-2 antibody. Middle panel: EC extracts were immunoprecipitated by incubated with protein-G Dynabeads® coupled to two different ITGA5 antibodies (as indicated). Immunoprecipitated complexes were subjected to Western blot analysis using antibodies against ITGA5 and NRP2 respectively. Right panel: NRP2 silencing was confirmed by subjecting the total cell lysate to Western blot analysis and incubating blots in antibodies against NRP-2 and HSC70. (**C**): Control siRNA-transfected ECs were prepared as described in **Fig2D** legend. Fixed ECs were incubated in primary antibodies against NRP2 and ITGA5. Images were captured using a Zeiss AxioImager M2 microscope at 63x magnification. Arrows indicate co-localisation of NRP2 and ITGA5. (**D**): siRNA-transfected ECs were prepared as described in **Fig2D** legend, however ECs were allowed to adhere overnight prior to fixation. ECs were incubated in primary antibody against ITGA5. (**E**) siRNA-transfected ECs were prepared as described in **D**, however fixed cells were incubated in ITGA5 primary antibody and phalloidin-364. Arrows indicate localisation of ITGA5 with actin filaments. Image panels shown in **C**-**E** are representative of n ≥10 cells per condition/treatment.

Studies have previously reported NRP1 to complex with ITGA5 in HUVECs [7] and NRP2 to complex with ITGA5 from co-cultures between HUVECs and renal cell carcinoma [42]. To investigate whether a direct interaction between NRP2 and ITGA5 exists in mLMECs, lysates were subjected to a series of co-immunoprecipitation studies followed by Western blot analysis. We found both NRP2 and ITGA5 co-immunoprecipitate with each other, indicating a physical interaction between the two receptors (**Fig3B**). Immuno-staining using a highly specific NRP2 antibody, reported previously not to cross-react with NRP1 [38, 45], also showed a strong co-localisation between NRP2 and ITGA5 close to the cell membrane (**Fig3C**). Given the significant upregulation of ITGA5 subunit expression following NRP2 depletion, and evidence to suggest a physical interaction between NRP2 and ITGA5, we next examined whether NRP2 depletion elicited an effect on ITGA5 localisation within our immortalised cells. Following treatment with either control or NRP2 siRNA, ECs were seeded overnight on a FN matrix and subsequently fixed to visualise endogenous ITGA5 expression. Compared to control siRNA treated ECs, NRP2 depleted ECs exhibited significant disruptions in ITGA5 organisation: ITGA5 appeared in elongated fibrillar-like structures (**Fig3D**), reminiscent of what has been described as fibrillar adhesions [28]. ImageJ^TM^ analysis of these ITGA5 containing structures confirmed a significant increase in the length of ITGA5 fibrils in NRP2 siRNA treated ECs suggestive of a disruption in ITGA5 trafficking (**Suppl. Fig3A**). We see this NRP2-dependent ITGA5 phenotype when using multiple NRP2 specific siRNAs (**Suppl. Fig3B**) and in primary ECs (**Suppl. Fig3C-D**). It is becoming increasingly clear that microtubule-actin highways co-ordinately regulate a range of intracellular trafficking mechanisms, including integrin transport [7, 46-49]. Co-immunostaining for both phalloidin and ITGA5 revealed that whilst ITGA5 localised to the ends of actin filaments in control treated ECs, at what we assume to be FAs, in NRP2 depleted cells, the elongated ITGA5 fibrils share a strong co-localisation along the actin filaments themselves (**Fig3E**). We believe this to support a mechanism of actin-dependent ITGA5 trafficking, whereby loss of NRP2 impedes the transport of ITGA5 to FAs at the leading edge of the cell.

### ITGA5 trafficking is dependent on NRP2 in ECs

Whilst a role for NRP2 in trafficking ITGA5 is novel, NRP1 has been shown previously to promote endocytosis of active α5β1 integrin through the Rab5 pathway [7]. In order to take an unbiased approach to elucidating candidate trafficking proteins NRP2 may associate with to regulate ITGA5 localisation in ECs, we used Label-Free quantitative mass spectrometry. Mass spectrometry was performed on two WT EC lines, transfected either with control or NRP2 siRNA, and immunoprecipitated for NRP2. siRNA-mediated depletion of NRP2 was confirmed for both cell lines. This analysis revealed proteins immunoprecipitating with NRP2 at a significantly increased fold-change compared to proteins analysed from a NRP2 knockdown cell-lysate. Shown are protein hits detected in both cell lines, including both ITGA5 and ITGB1. A number of endocytic trafficking proteins were also detected as candidate binding partners of NRP2, including clathrin, caveolin-1, lamtor1, scamp1, and annexin-A1 (**Fig4A**).

**Figure 4.**
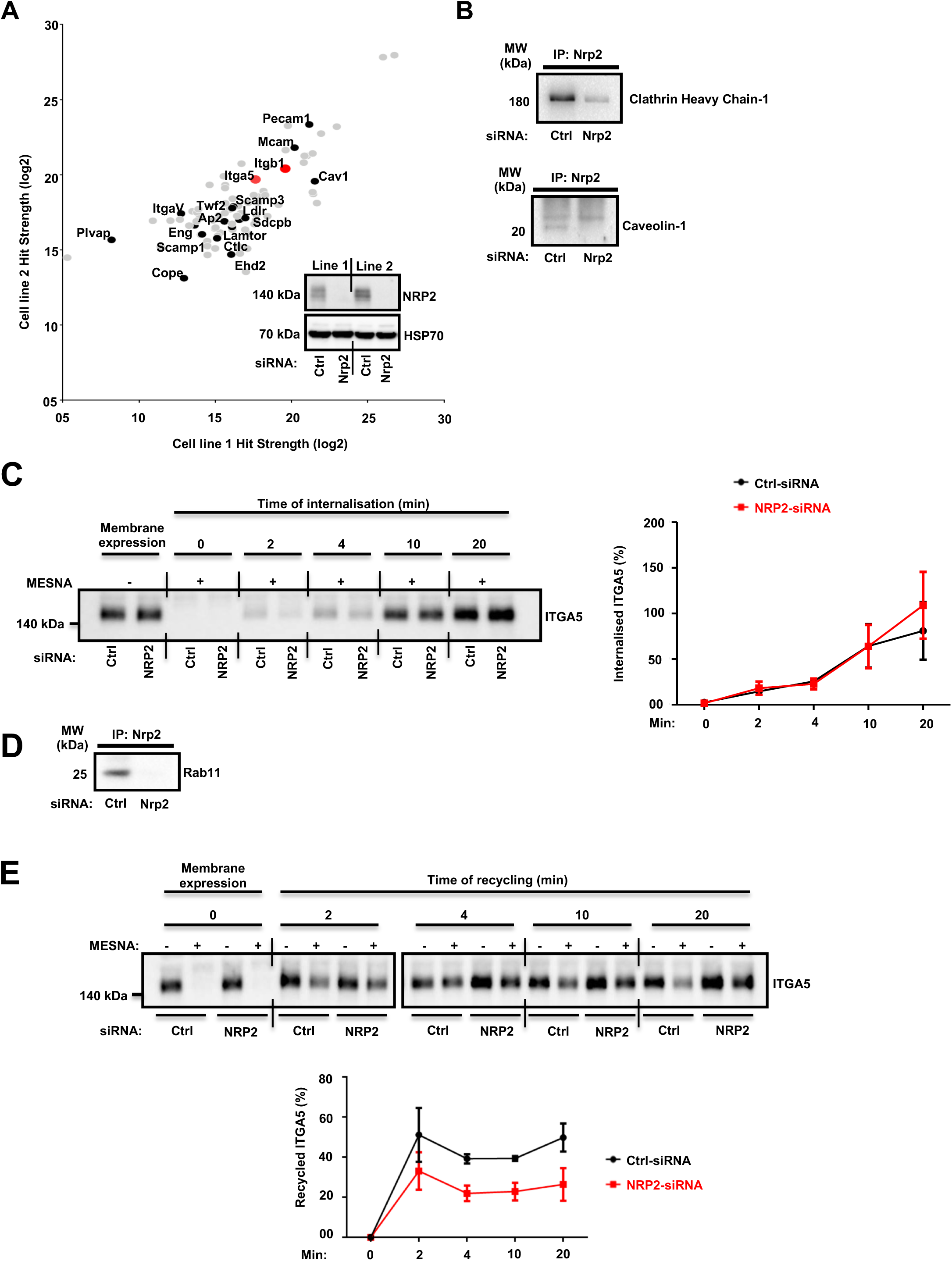
ITGA5 trafficking in ECs is NRP2 dependent. (**A**): Label-free quantitative mass spectrometry peptide hits identified using MaxQuant software from the Andromeda peptide database. Panel depicts peptide hits detected at a higher fold-change in two control siRNA treated EC lines than those detected from two NRP2 siRNA treated EC lines. NRP2 silencing was confirmed by subjecting EC extracts to Western blot analysis. (**B**): siRNA-transfected ECs were seeded onto FN and incubated for 48 hours. EC extracts were immunoprecipitated by incubation with protein-G Dynabeads® coupled to a NRP2 antibody. Immunoprecipitated complexes were subjected to Western blot analysis using antibodies against clathrin heavy chain-1 (top) and caveolin-1 (bottom). (**C**): siRNA-transfected ECs were seeded onto FN and incubated for 48 hours. ECs were subsequently starved in serum-free media, before being placed on ice. EC cell surface proteins were labelled with 0.3 mg/ml biotin. ECs were then incubated for the indicated timepoints at 37°C, 5% CO_2_. A sample of ECs were maintained at 4°C for use as positive/negative (+/-Mesna) controls. Following incubation at 37°C, ECs were placed on ice and incubated with 100 mM Mesna. EC lysates were then immunoprecipitated with protein-G Dynabeads® coupled to an anti-biotin antibody. Immunoprecipitated biotin-labelled proteins were separated by SDS-PAGE and subjected to Western bolt analysis. The level of internalised ITGA5 at each time of incubation was normalised to the (-) Mesna control. Left panel shows Western blot image, right panel shows the mean densitometric analysis obtained using ImageJ^TM^. Also shown is confirmation of NRP-2 silencing. N=3 independent experiments. (**D**): Co-immunoprecipitation study as described in **B**. Immunoprecipitated complexes were subjected to Western blot analysis using an anti-Rab11 antibody. (**E**): siRNA-transfected ECs were prepared as described in **C**. After biotin surface labelling, ECs were incubated in serum free media for 20 minutes at 37°C to allow for internalisation. A sample of ECs were maintained at 4°C for use as positive/negative controls. The remaining ECs were then placed on ice, and any un-internalised biotin-labelled proteins stripped off using 100 mM Mesna. The internalised protein fraction was allowed to recycle by incubating the ECs for the indicated timepoints at 37°C. ECs were then returned to ice and incubated in 100 mM Mesna. No Mesna treatment dishes at each timepoint were used as comparative controls. All subsequent stages were performed in the same manner as described in **C**. The level of the recycled ITGA5 was determined by normalising the amount of ITGA5 quantified from dishes treated with Mesna to the total ITGA5 on the membranes of the Mesna-untreated cells in the same period of incubation. N=3 independent experiments.

α5β1 integrin is known to be internalised both by clathrin and caveolae-mediated endocytosis [50, 51]. In order to validate a potential mechanism by which NRP2 regulates ITGA5 internalisation via its contact with clathrin and caveolin, we performed coimmunoprecipitation assays to prove a physical interaction. NRP2 coimmunoprecipitated with both clathrin heavy chain-1 and caveolin-1 (**Fig4B**), supporting the mass spectrometry results. We subsequently conducted cell surface biotinylation assays to examine ITGA5 internalisation directly, in ECs treated either with control siRNA or NRP2 siRNA. We observed no change in the rate of ITGA5 internalisation in NRP2 depleted ECs (**Fig4C**), suggesting that NRP2 does not regulate internalisation of total ITGA5 levels. We subsequently considered whether NRP2 regulates ITGA5 recycling back to the membrane.

Integrin recycling is mediated by Rab GTPase proteins [52, 53], specifically, α5β1 integrin follows long-loop recycling via a Rab11 dependent mechanism [54, 55]. In addition to investigating whether an interaction between NRP2 and Rab11 exists, we also considered interactions with Rab4 and Rab5, proteins both involved in regulating short-loop recycling [50]. NRP2 coimmunoprecipitated with Rab11 only (**Fig4D**), and not Rab4, or Rab5 (data not shown). Importantly, NRP2 did not co-immunoprecipitate with NRP1 in any of our studies (data not shown), suggesting these two structurally related proteins regulate ITGA5 trafficking through distinct pathways. As our mass spectrometry analysis had identified candidate interactions between NRP2 and other trafficking and recycling molecules, such as Rab11Fip5, an adaptor for Rab11 recycling vesicles and previously reported to co-immunoprecipitate with ITGA5, we performed biotin recycling assays. In these assays, the biotin-labelled cell surface proteins were allowed to internalise before stripping off any remaining surface biotin. Cells were then incubated over 20 minutes to stimulate the recycling process. In contrast to our internalisation assays, NRP2 silencing significantly attenuated the rate of total ITGA5 recycling back to the membrane compared to ECs treated with control siRNA (**Fig4E**). We therefore present a potential mechanism whereby NRP2 promotes mLMEC migration and adhesion to FN by regulating actin dependent ITGA5 recycling back to the cell membrane.

## Discussion

NRP2 is known to regulate ECM adhesion and migration in various cell lines, including human ECs [21], immune cells [56] and cancer cells[15, 20, 26, 42, 57]. In the latter, it has been described as a biomarker for poor prognosis because its expression correlates with increased migratory and invasive behaviour [15, 26, 42, 57]. In many of these examples, NRP2 promotes cell adhesion and migration by exerting influence over VEGF-mediated pro-angiogenic pathways and inducing integrin engagement. ITGA5 and FN are known to be upregulated in ECs during angiogenesis [40, 41], and are key players in formation of neo-vasculature [58, 59]. The data we present here support a model whereby NRP2 promotes EC adhesion and migration on FN by regulating actin-dependent ITGA5 recycling. Importantly, the means by which it does so appears to be mechanistically distinct from that of NRP1, which is known to regulate internalisation of active ITGA5 via a rab5-dependent pathway [7].

We previously demonstrated an ITGB3-dependent role for NRP1 in mediating EC migration and FA turnover on FN. EC migration on FN becomes NRP1 dependent when ITGB3 levels are perturbed [8]. Like NRP1, NRP2 expression is upregulated in ITGB3 depleted cells (**Fig1A**). Given this observation, and the structural and domain homologies between NRP1 and NRP2 [3], we sought to investigate whether NRP2 shared a similar regulatory interplay with ITGB3. We demonstrated by siRNA knockdown that EC migration depends on NRP2, but this is independent of ITGB3 expression levels (**Fig1C**).

Like NRP1, NRP2 also regulates a number of integrin-dependent cellular processes on FN. Upon NRP2 knockdown, EC adhesion is reduced (**Fig2A**), FA dynamics are altered (**Fig2C-D**), and Rac1 activation is impaired (**Fig2E**) [7, 8]. Rac1 was also identified from our mass spectrometry analysis as a candidate binding partner of NRP2, which supports these findings. Integrins provide the structural link that allows for: (1) adhesion to the ECM; (2) anchorage of actin stress fibres to the membrane; (3) the generation of force that is required for migration [54, 60, 61]. Both αvβ3- and α5β1-integrins are FN receptors in ECs [58, 59], but given the lack of any ITGB3 input into the NRP2 processes we were examining, and the upregulation of ITGA5 expression upon NRP2 depletion (**Fig3A**), we focussed our attention on this integrin subunit. Cao *et al*. [42] demonstrated NRP2 to interact with ITGA5 in co-cultures between HUVECs and renal cell carcinoma to promote FN-mediated adhesion. To our knowledge, though, we are the first to show evidence of a direct interaction between NRP2 and ITGA5 in microvascular ECs (**Fig3B-C**). Furthermore, our findings demonstrate that NRP2 modulates ITGA5 function by regulating its subcellular trafficking along the actin cytoskeleton. Importantly, the disrupted ITGA5 phenotype we see following NRP2 depletion in our immortalised ECs is also present in primary ECs. Along with our previous studies demonstrating they express a canonical adhesome [32], this rebuts the concept that immortalised ECs are an inappropriate model for studying angiogenic processes.

NRP2’s role in trafficking ITGA5 in ECs also appears to be distinct from that of NRP1. It has been reported that NRP1 regulates active α5β1 integrin endocytosis via GIPC1 [7], however interactions between GIPC and NRP2 were not detected from either mass spectrometry analysis or co-immunoprecipitation studies (data not shown). Although we did not observe any changes in internalisation of total ITGA5 upon NRP2 depletion (**Fig4C**), we were unable to examine trafficking of active α5β1 in our murine cells. In fact, we observed physical interactions between NRP2 and both clathrin and caveolin by immunoprecipitation (**Fig4B**) suggesting NRP2 at least has the potential to regulate the endocytosis of active α5β1 integrin. Following internalisation by clathrin or caveolin-dependent mechanisms, α5β1 integrin has been shown to undergo long-loop recycling back to the membrane within Rab11 positive vesicles [54, 55]. To our knowledge neither NRP1 or NRP2 have been reported to regulate ITGA5 recycling in microvascular ECs. Not only do we show NRP2 depletion slows total ITGA5 recycling (**Fig4D**), we also show NRP2 co-immunoprecipitates with Rab11 in our ECs (**Fig4E**).

Finally, the elevated total cellular levels of ITGA5 that occur when NRP2 is knocked-down, particularly in light of no changes at the cell surface (data not shown), suggest other points in the ITGA5 life cycle are governed by NRP2. Given NRP2’s interactions with clathrin and caveolin, both known to play a role in endosomal trafficking to lysosomes [50, 51, 62], we speculate a role for the molecule in regulating ITGA5 degradation. We support this hypothesis by showing NRP2 also immunoprecipitates with both Lamtor1 and Scamp1 (**Fig4A**), proteins previously shown to regulate lysosomal trafficking [63, 64].

In close, we propose a novel mechanism by which NRP2, independently of its role as a coreceptor for VEGF-A, promotes Rac1-mediated EC adhesion and migration to FN matrices by regulating recycling of ITGA5. Importantly, we provide evidence to suggest that NRP2 acts in a mechanistically distinct manner to NRP1. Finally, whilst we have not yet shown any phenotypic interactions between NRP2 and ITGB3, we cannot rule out a role for ITGB3 in regulating NRP2 trafficking. Like NRP1, NRP2 levels are significantly elevated when ITGB3 expression is reduced (**Fig1A**). Our findings allude to a complex interplay between FN-binding integrins and neuropilins which regulate EC migration.

## Materials and Methods

### Animal Generation

All experiments were performed in accordance with UK home office regulations and the European Legal Framework for the Protection of Animals used for Scientific Purposes (European Directive 86/609/EEC), prior to the start of this project. Transgenic mice expressing a knockout for the β3-integrin allele (KO-Itgb3) were generated by substituting a 1.4 kb HindIII fragment of the β3 gene including exons I and II with a 1.7 kb construct containing a Pgk-neomycin (neo)-resistance cassette (**Suppl. Fig1G**). The PCR analysis was carried out using the following oligonucleotide primers as previously described by Hodivala-Dilke *et al*. [65]. Forward primer 1: 5’-CTTAGACACCTGCTACGGGC-3’ Reverse primer 2: 5’-CACGAGACTAGTGAGACGTG-3’.

### Cell Isolation, Immortalisation and Cell Culture

Primary mouse lung microvascular endothelial cells (mLMECs) were isolated from adult mice bred on a mixed C57BL6/129 background. Primary ECs were twice positively selected for their expression of intracellular adhesion molecule-2 (ICAM-2) by magnetic activated cell sorting (MACS) as previously described by Reynolds & Hodivala-Dilke [66]. ECs were immortalised using polyoma-middle-T-antigen (PyMT) retroviral transfection as previously described by Robinson *et al*. [23]. Immortalised mLMECs were cultured in IMMLEC media, a 1:1 mix of Ham’s F-12:DMEM medium (low glucose) supplemented with 10% FBS, 100 units/mL penicillin/streptomycin (P/S), 2 mM glutamax, 50 μg/mL heparin (Sigma). Immortalised mLMECs were cultured on 0.1% gelatin coated flasks at 37°C in a humidified incubator with 5% CO_2_. For experimental analyses, plates, dishes, flasks and coverslips were coated in 10 µg/ml human plasma fibronectin (FN) (Millipore) overnight at 4°C. Vascular endothelial growth factor-A (VEGF-A164: mouse equivalent of VEGF-A165) was made in-house as previously described by Krilleke *et al*. [67].

### siRNA Transfection

ECs were transfected with non-targeting control siRNA or a mouse-specific NRP2 siRNA construct (Dharmacon), suspended in nucleofection buffer (200 mM Hepes, 137 mM NaCl, 5 mM KCl, 6 mM D-glucose, and 7 mM Na2HPO4 in nuclease-free water) using either the Amaxa nucleofector system II (Lonza) under nucleofection program T-005 or the Amaxa 4Dnucleofector system (Lonza) under nucleofection program EO-100 according to manufacturer’s instructions.

### Western Blot Analysis

siRNA transfected ECs were seeded into FN-coated 6-well plates at a seeding density of 5×10^5^ cells/well and incubated for 48 hours at 37°C in a 5% CO_2_ incubator. ECs were lysed in electrophoresis sample buffer (ESB) (Tris-HCL: 65 mM pH 7.4, sucrose: 60 mM, 3% SDS), and homogenised using a Tissue Lyser (Qiagen) with acid-washed glass beads (Sigma). Following protein quantification using the DC BioRad assay, 30 µg of protein from each sample was loaded onto 8% polyacrylamide gels and subjected to SDS-PAGE. Proteins were transferred to a nitrocellulose membrane (Sigma) and incubated in 5% milk powder in PBS 0.1% Tween-20 (0.1% PBST) for 1 hour at room temperature followed by an overnight incubation in primary antibody diluted 1:1000 in 5% bovine serum albumin (BSA) in 0.1% PBST at 4°C. Membranes were washed 3x with 0.1% PBST and incubated in an appropriate horseradish peroxidase (HRP)-conjugated secondary antibody (Dako) diluted 1:2000 in 5% milk powder in 0.1% PBST for 2 hours at room temperature. Membranes were washed again 3x with 0.1% PBST before being incubated with Pierce ECL Western Blotting Substrate solution (Thermo Scientific). Chemiluminescence was detected on a ChemiDoc^TM^ MP Imaging System darkroom (BioRad). Densitometric readings of band intensities for blots were obtained using ImageJ^TM^. Primary antibodies (all used at 1:1000 dilution and purchased from Cell Signalling Technology, unless noted otherwise) were: anti-NRP2 (clone D39A5), anti-ITGB3 (clone 4702S), anti-HSC70 (clone B-6, Santa Cruz Biotechnology), anti-phospho VEGFR2 (Y1175) (clone 2478), anti-VEGFR2 (clone 2479), anti-NRP1 (clone 3725S), anti-phospho ERK1/2 (clone 9101), anti-ERK1/2 (clone 4695), anti-ITGA5 (clone 4705S), anti-ITGB1 (clone ab179471, Abcam), anti-clathrin heavy chain-1 (clone ab21679, Abcam), anti-caveolin-1 (clone ab18199, Abcam), anti-Rab11 (clone 3539), ERG: Ab92513, Pecam-1 (77699), prox-1 (clone ab11941, Abcam), claudin-5 (clone ab131259, Abcam), VE-cadherin (clone ab205336, Abcam).

### Signalling Assays

siRNA-transfected ECs were seeded into FN-coated 6 cm cultures dishes at a density of 5×10^5^ cells/well and incubated for 48 hours at 37°C and 5% CO_2_. ECs were then PBS washed and starved for 3 hours in serum free medium (OptiMEM®; Invitrogen). VEGF was then added at a final concentration of 30 ng/ml. After the desired time of VEGF-stimulation, ECs were subjected to lysing, protein quantification and protein expression analysis by Western blot.

### Random Migration Assays

siRNA-transfected ECs were seeded into FN coated 24-well plates at a density of 7×10^4^ cells/well 24 hours post nucleofection, and allowed to adhere for 3 hours at 37°C and 5% CO_2_. Fixed images of multiple fields/well were taken every 10 minutes for 15 hours at 37°C and 5% CO_2_ using an inverted Zeiss Axiovert microscope with one-phase contrast. Random migration was quantified by manually tracking individual cells using the ImageJ^TM^ plugin mTrackJ. Random migration speed was calculated in µm/hour.

### Colorimetric Adhesion Assays

siRNA-transfected ECs were seeded into 96-well plates at a density of 4×10^4^ cells/well 48 hours post nucleofection. 96-well plates were pre-coated with FN overnight, then blocked in 5% BSA for 1 hour at room temperature. ECs were then incubated at 37°C in a 5% CO_2_ incubator for the indicated timepoints, in addition to a 3-hour incubation control plate. Following incubation, ECs were washed 3x with PBS + 1 mM MgCl2 + 1 mM CaCl2, fixed in 4% PFA, and stained with methylene blue for 30 minutes at room temperature. ECs were washed in dH_2_O and air-dried, before the dye from stained adhered ECs was extracted by a de-stain solution (50% ethanol, 50% 0.1M HCL). The absorbance of each well was then read at 630 nm. Data was normalised to the relative number of the total cells seeded in the 3-hour incubation plate.

### Focal Adhesion Turnover Assays

ECs double transfected with control or NRP2 siRNA and a GFP-tagged paxillin construct (kindly provided by Dr Maddy Parsons, Kings College, London) were seeded onto FN-coated acid-washed, oven sterilised glass coverslips in 24-well plates at a seeding density of 4×10^4^ cells/well. 48 hours post nucleofection, coverslips were PBS washed, fixed in a Ludin chamber (Life Imaging Services GmbH), and live imaged in OptiMEM® phenol-red free medium supplemented with 2% FBS and P/S at 37°C and 5% CO_2_ using an inverted Axiovert (Carl Zeiss Ltd) microscope in which an individual cell was captured every one minute for 30 minutes. Focal adhesion (FA) assembly and disassembly speeds were analysed by manually tracking the number of selected GFP-paxillin-positive focal adhesions using the Image J^TM^ MTrackJ plugin software.

### Immunocytochemistry

siRNA-transfected ECs were seeded onto FN-coated acid-washed, oven sterilised glass coverslips in 24-well plates at a seeding density of 2.5×104 cells/well, and incubated at 37°C and 5% CO_2_. ECs were fixed at indicated timepoints in 4% paraformaldehyde (PFA) for 10 minutes, washed in PBS, blocked and permeabilised with 10% goat serum, PBS 0.3% triton X-100 for 1 hour at room temperature. Cells were incubated in primary antibody diluted 1:100 in PBS overnight at 4°C. Primary antibodies were: anti-paxillin (clone ab32084; Abcam), anti-ITGA5 (clone ab150361; Abcam). Coverslips were PBS washed, and incubated with donkey anti-rabbit Alexa fluor-488 secondary antibody diluted 1:200 in PBS for 2 hours at room temperature. F actin staining was performed by incubating cells in phalloidin-364 diluted 1:40 in PBS for 2 hours at room temperature during secondary antibody incubation. Coverslips were PBS washed again, before being mounted onto slides with Prolong® Gold containing DAPI (Invitrogen). Images were captured using a Zeiss AxioImager M2 microscope (AxioCam MRm camera) at 63x magnification. FA number and size was quantified using ImageJ^TM^ software as previously described by Lambert *et al*. [68]. A FA size lower detection limit was set at 0.8 microns. ITGA5 length was measured using the ImageJ^TM^ software plugin simple neurite tracer.

### Rac1 Pulldown

siRNA-transfected ECs were seeded onto FN at a density of 2×10^5^ cells/dish and incubated for 180 minutes at 37°C and 5% CO_2_. ECs were lysed on ice with MLB (Sigma) (diluted to 1X with sterile water containing 10% glycerol and 1X Halt^TM^ protease inhibitor cocktail (Thermo Scientific). EC lysates were cleared by centrifugation. EC extracts were immunoprecipitated by incubation with 10 µg Rac1 assay reagent (PAK-1 PBD magnetic beads, Sigma) for 45 minutes at 4°C with gentle agitation. Immunoprecipitated complexes were subjected to Western blot analysis. Nitrocellulose membranes were immunoblotted using 1 µg/mL of anti-Rac1, clone 23A8 (Sigma).

### Co-Immunoprecipitation Assays

siRNA-transfected ECs were seeded into FN-coated 10 cm dishes at a density of 2×10^6^ cells/dish, and incubated for 48 hours at 37°C and 5% CO_2_. ECs were then lysed on ice in lysis buffer as previously described by Valdembri *et al.* [7] in the presence of 1X Halt protease inhibitor cocktail (Thermo Scientific) and protein quantified using the DC BioRad assay. 100 µg protein from each sample was immunoprecipitated by incubation with protein-G Dynabeads® (Invitrogen) coupled to a rabbit anti-NRP2 antibody (clone 3366, Cell Signalling Technology) on a rotator overnight at 4°C. Immunoprecipitated complexes were then washed 3x with lysis buffer + 1X Halt^TM^ protease inhibitor cocktail, and once in PBS, before being added to and boiled in NuPAGE sample reducing agent and sample buffer (Life Technologies) for Western blot analysis.

### Co-localisation Assays

siRNA-transfected ECs were prepared the same as for immuno-cytochemistry. Cells were incubated in primary antibody diluted 1:50 in PBS overnight at 4°C. Primary antibodies were: anti-NRP2 (clone sc-13117, Santa Cruz Biotechnology), and anti-ITGA5 (clone ab150361; Abcam). Coverslips were PBS washed, and incubated with both donkey anti-rabbit Alexa fluor-488, and goat anti-mouse Alexa fluor-546 secondary antibodies diluted 1:200 in PBS for 2 hours at room temperature. Coverslips were PBS washed again, before being mounted onto slides with Prolong® Gold containing DAPI (Invitrogen). Images were captured using a Zeiss AxioImager M2 microscope (AxioCam MRm camera) at 63x magnification.

### Mass Spectrometry Analysis

NRP2 co-immunoprecipitation samples were sent to Fingerprints Proteomics Facility (Dundee University, UK), which carried out label-free quantitative mass spectrometry and peptide identification using the MaxQuant software based on the Andromeda peptide database as described by Schiller *et al*. [69]. Fig4A depicts peptide hits detected at a higher fold-change in two control siRNA treated EC lines than those detected from two NRP2 siRNA treated EC lines.

## Internalisation and Recycling Assays

### Internalisation

siRNA-transfected ECs were seeded into FN-coated 10 cm dishes at a density of 2×10^6^ cells/dish, and incubated for 48 hours at 37°C and 5% CO_2_. ECs were then starved in serum-free OptiMEM® for 3 hours at 37°C in a 5% CO^2^ incubator, before being placed on ice for 5 minutes, then washed twice with Soerensen buffer (SBS) pH 7.8 (14.7mM KH2PO4, 2mM Na2HPO4, and 120mM Sorbitol pH 7.8) as previously described by Remacle et al. [70]. EC cell surface proteins were labelled with 0.3 mg/ml biotin (Thermo Scientific) in SBS for 30 minutes at 4°C. Unreacted biotin was quenched with 100 mM glycine for 10 minutes at 4°C. ECs were then incubated in pre-warmed serum-free OptiMEM® for the indicated time points at 37°C in a 5% CO^2^ incubator. A sample of ECs were maintained at 4°C for use as positive/negative (+/-Mesna) controls. Following incubation, dishes were immediately placed on ice, washed twice with SBS pH 8.2, and incubated with 100mM Mesna (Sigma) for 75 minutes at 4°C (with the exception of Mesna control plates, which were lysed in lysis buffer (25mM Tris-HCl, pH 7.4, 100mM NaCl, 2mM MgCl2, 1mM Na3VO4, 0.5 mM EGTA, 1% Triton X-100, 5% glycerol, and protease inhibitors), and placed on ice). Following Mesna incubation, excess Mesna was quenched with 100mM iodoacetamide (Sigma) for 10 minutes at 4°C, then ECs were washed twice with SBS pH 8.2 and lysed. Lysates were cleared by centrifugation at 12,000g for 20 minutes at 4°C. Supernatant proteins were then quantified using the DC BioRad assay, and subsequently immunoprecipitated with Dynabeads coupled to mouse anti-biotin antibody overnight at 4°C. Immunoprecipitated biotin-labelled proteins were separated by SDS-PAGE and subjected to Western bolt analysis. The level of internalised ITGA5 at each time of incubation was normalised to the (-Mesna) control.

### Recycling

After surface labelling, ECs were incubated in pre-warmed serum free OptiMEM® for 20 minutes at 37°C to allow internalisation. A sample of ECs were maintained at 4°C for use as positive/negative controls. The remaining dishes were subsequently placed on ice, washed twice with SBS pH 8.2, and any un-internalised biotin-labelled proteins were stripped off using 100 mM Mesna in Tris buffer for 75 minutes at 4°C. The internalised fraction of proteins was then allowed to recycle to the membrane by incubating the ECs for the indicated time points in serum-free OptiMEM® at 37°C. Following the indicated times of incubation, dishes were placed on ice, washed twice with SBS pH 8.2, and subjected to Mesna incubation for 75 minutes at 4°C. No Mesna treatment dishes at each timepoint were used as controls. All subsequent stages were performed in the same manner as the internalisation assay. The level of the recycled ITGA5 was determined by normalising the amount of ITGA5 quantified from dishes treated with Mesna, to the total ITGA5 on the membranes of the Mesnauntreated cells in the same period of incubation.

### Statistical Analysis

The graphic illustrations and analyses to determine statistical significance were generated using GraphPad Prism 6 software and Student’s t-tests, respectively. Bar charts show mean values and the standard error of the mean (±SEM). Asterisks indicate the statistical significance of P values: P > 0.05 = ns (not significant), * P < 0.05, ** P < 0.01, *** P < 0.001 and **** P < 0.0001.

## Acknowledgments

This work was supported by funding from: the UKRI Biotechnology and Biological Sciences Research Council Norwich Research Park Biosciences Doctoral Training Partnership (Grant numbers BB/M011216/1, BB/J014524/1); BHF (grant number PG/15/25/31369); The Ministry of Education, Saudi Arabia. Additionally, we thank Norfolk Fundraisers, Mrs Margaret Doggett, and the Colin Wright Fund for their kind support and fundraising over the years. Robinson is also partially funded by the BBSRC Institute Strategic Programme Gut Microbes and Health BB/R012490/1 and its constituent project (BBS/E/F/000PR10355).

## Supplementary Figure Legends

**Suppl. Fig1.**
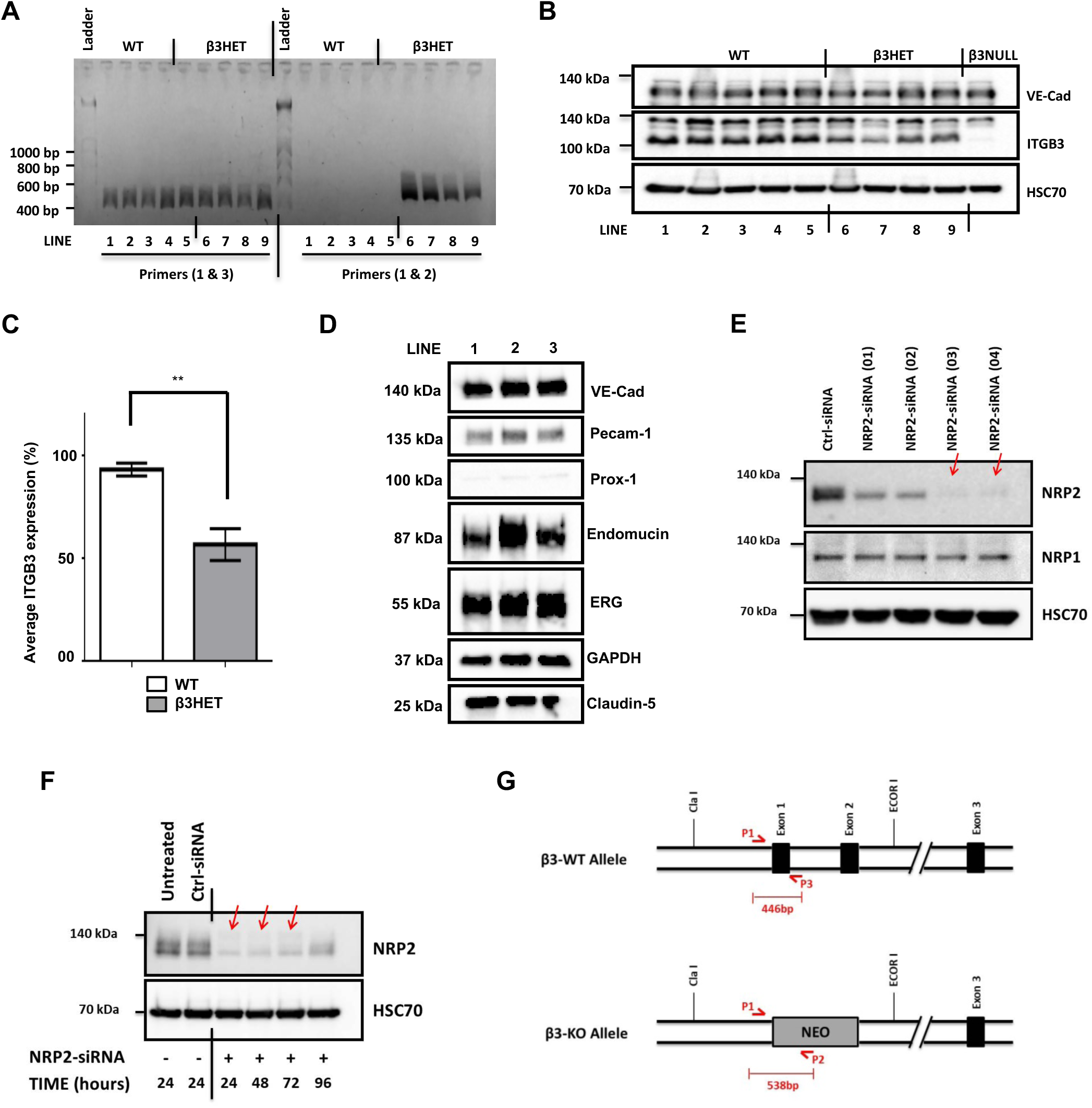
(**A**): PyMT transfected mLMEC pellets from 9 different immortalised EC lines were subjected to DNA extraction and PCR-based genotyping at the ITGB3 locus. Shown is an agarose gel of P1/P3 and P1/P2 primer products from all EC lines. Lines 1-5 show only a P1/P3 product, and are therefore wild-type (WT) for the ITGB3 locus, whilst lines 6-9 show both a P1/P3 product, and a P1/P2 product, and are therefore heterozygous (β3HET) for the ITGB3 locus. (**B**): Western blot analysis of VE-Cadherin and ITGB3 expression in the same clones shown in **A**. HSC70 was used as a loading control. Because the antibody used for detection of ITGB3 recognises a non-specific band at approximately (135kDa), a β3-knockout (NULL) lysate was included as a control (cell line #10). (**C**): Left panel shows the densitometric analysis of mean ITGB3 band intensities normalised against HSC70 obtained using ImageJ^TM^ (from the Western blot shown in **B**). ** P<0.01. (**D**): The EC identity of 3 immortalised EC lines was re-confirmed by Western blot analysis, immunoblotting against additional EC markers Pecam-1, Endomucin, ERG and Claudin-5, alongside a GAPDH loading control and a lymphatic marker Prox-1. (**E**): ECs were transfected either with control siRNA or one of four different NRP2-specific siRNAs (01 - 04) and incubated for 48 hours. EC extracts were then subjected to Western blot analysis using antibodies against NRP2, NRP1 and HSC70. Except where noted (Suppl. Figure 3), NRP2 siRNA #03 was used for all subsequent experiments to silence NRP2 expression. (**F**): siRNA-transfected ECs were incubated for the indicated timepoints before being lysed and subjected to Western blot analysis using antibodies against NRP2 and HSC70. Asterisks indicate statistical significance from unpaired two-tailed t-tests. (**G**): Schematic diagram of the ITGB3 (WT) and (HET) loci, showing where PCR genotyping primers align. P1 and P3 amplify a wild-type product of 446-bp, whilst P1 and P2 amplify a knockout product of 538-bp.

**Suppl. Fig2.**
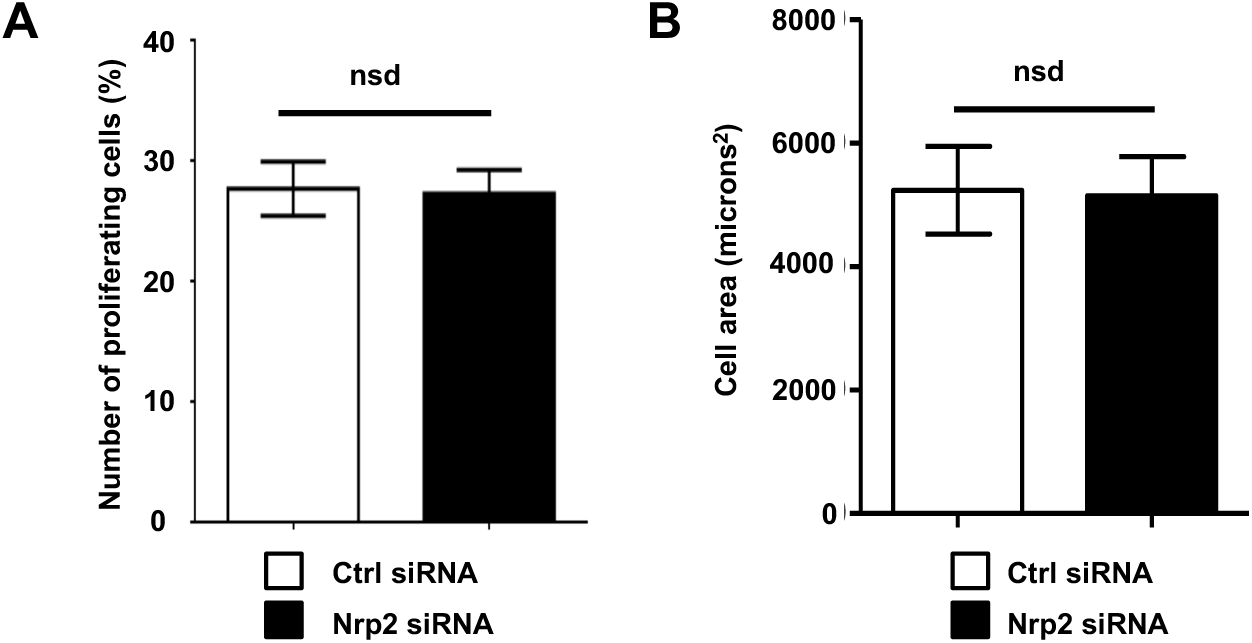
(**A**): siRNA-transfected ECs were seeded onto FN and incubated for 30 hours. ECs were then re-seeded onto FN-coated coverslips, and allowed to adhere for 4 hours in serum-free media. Media was then replaced with 10 µM BrdU in complete culture medium, and the cells incubated for 12 hours at 37°C. ECs were then fixed with 4% PFA, before hydrolysing the DNA with 1M HCL. ECs were permeabilised and blocked with Dako^®^ Protein Block Serum-Free. ECs were then incubated with anti-BrdU at 4°C overnight in a humidified chamber, followed by incubation in secondary antibody. Coverslips were subsequently mounted and the number of cells in S-phase (proliferating cells) was determined by dividing the number of the BrdU-labelled cells by the number of DAPI-labelled cells. n=19 independent fields of view, containing on average 50 cells per field, per condition. (**B**): Accompanying analysis to **Fig2D**. The cell area (microns^2^) was measured using ImageJ^TM^. Quantification performed on mean data from n≥25 ECs over N=3 independent experiments, nsd = not significantly different from unpaired two-tailed t-test.

**Suppl. Fig3.**
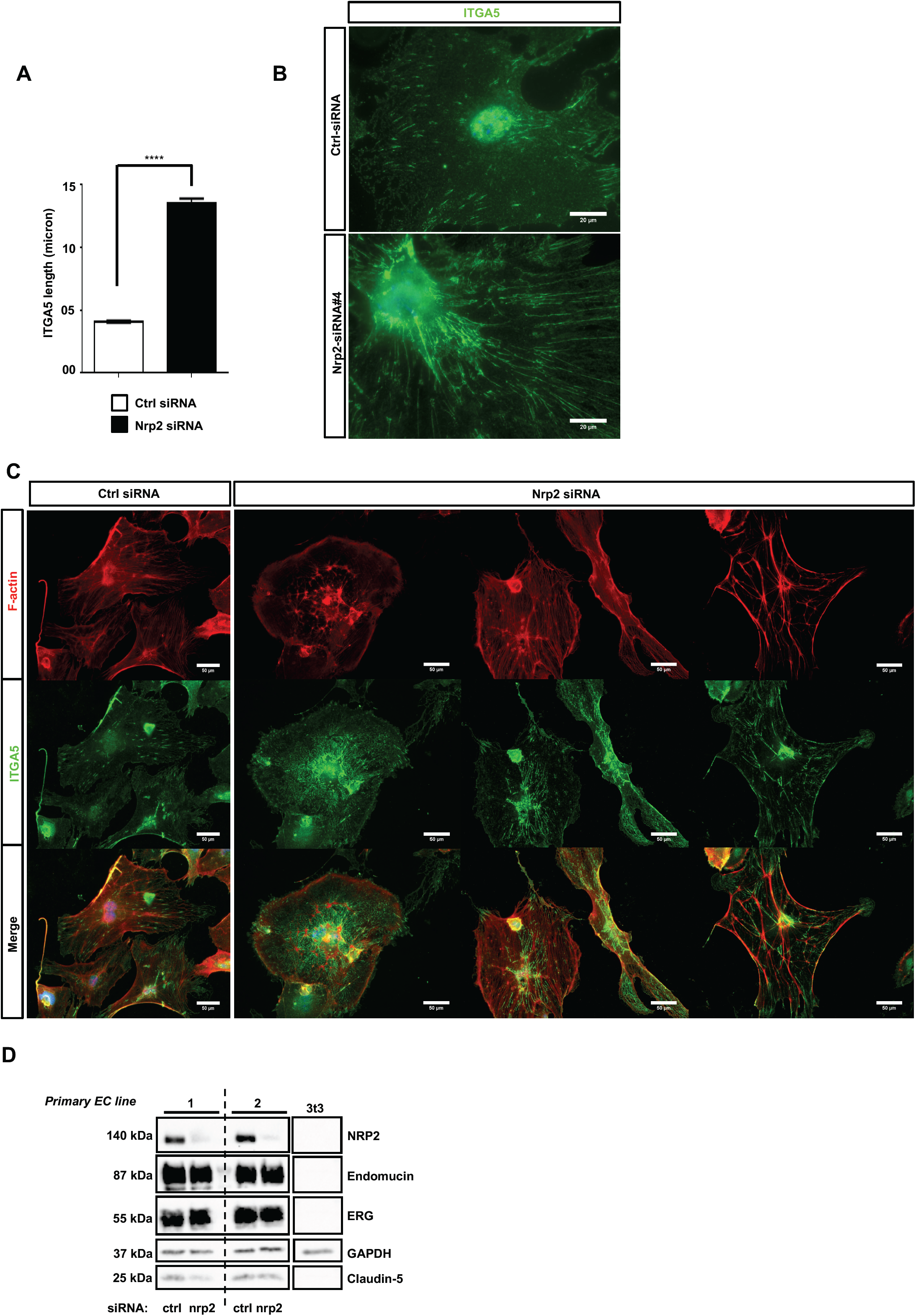
(**A**): Accompanying analysis to **Fig3D**. siRNA-transfected ECs were prepared for immunocytochemistry as described in **Fig2C**. Fixed ECs were incubated in primary antibodies against ITGA5 overnight at 4°C. ITGA5 length was measured using the ImageJ^TM^ software plugin simple neurite tracer. n≥490 cells per condition, ****P < 0.0001. Asterisks indicate statistical significance from unpaired two-tailed t-tests. (**B**): siRNA-transfected ECs from 3 different immortalised EC lines were prepared as described in **Fig3D** legend, however ECs were transfected with either control (top) or NRP2 siRNA#04 (bottom). Panels show representative images from N=3 independent lines, n≥10 cells per line. (**C**): Primary ECs were transfected with either ctrl or NRP2 siRNA and prepared for immunostaining as described in **Fig3E**. Panels show representative images from n≥10 cells per condition from 2 independent primary EC lines. (**D**): Western blot analysis of cell lysates from both primary EC clones alongside a lysate from a known fibroblast control cell line. EC extracts were immunoblotted using antibodies against known EC markers Endomucin, ERG and Claudin-5, alongside a GAPDH loading control.

**Suppl. Fig4.**
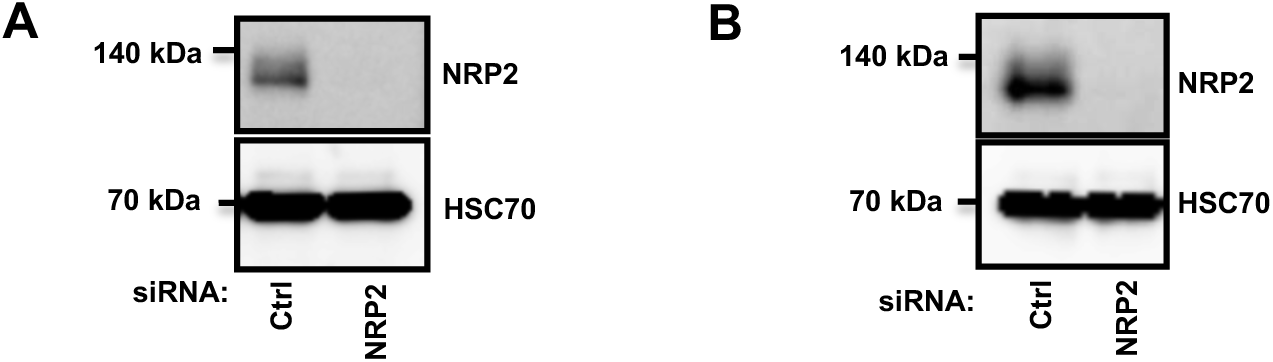
(**A**): Accompanying NRP2 knockdown confirmation for **Fig4C**. EC extracts were immunoblotted using antibodies to NRP2 and HSC70. (**B**): Accompanying NRP2 knockdown confirmation for **Fig4E**. EC extracts were immunoblotted using antibodies to NRP2 and HSC70.

